# A general approach to engineer positive-going eFRET voltage indicators

**DOI:** 10.1101/690925

**Authors:** Ahmed S. Abdelfattah, Rosario Valenti, Allan Wong, Minoru Koyama, Douglas S. Kim, Eric R. Schreiter

## Abstract

We engineered electrochromic fluorescence resonance energy transfer (eFRET) genetically encoded voltage indicators (GEVIs) with “positive-going” fluorescence response to membrane depolarization through rational manipulation of the native proton transport pathway in microbial rhodopsins. We transformed the state-of-the-art eFRET GEVI Voltron into Positron, with kinetics and sensitivity equivalent to Voltron but flipped fluorescence signal polarity. We further applied this general approach to GEVIs containing different voltage sensitive rhodopsin domains and various fluorescent dye and fluorescent protein reporters.

---

Genetically encoded voltage indicators (GEVIs) allow visualization of fast action potentials and subthreshold dynamics in groups of genetically defined neurons with high spatiotemporal resolution^1,2^. Monitoring voltage signals *in vivo* in large populations of neurons will enable dissection of detailed mechanistic links between brain activity and animal behavior. Despite recent advances, no current GEVI has ideal properties for routine, robust *in vivo* imaging. Knowledge of the mechanistic function of GEVIs will lead to improved designs. Existing GEVIs use voltage sensitive protein domains from voltage sensitive ion channels or phosphatases^3–7^ (VSD GEVIs), or microbial rhodopsin domains^8–14^ (rhodopsin GEVIs). Archaerhodopsin 3 (Arch) from *Halorubrum sodomense* was the first rhodopsin GEVI that accurately tracked changes in neuronal membrane potential^8^. However, Arch also generated a hyperpolarizing light-driven current upon exposure to imaging light^8,15^ by functioning as an outward proton pump^15^. A single amino acid substitution (D95N) in Arch abolished light-driven currents and retained Arch voltage-dependent fluorescence change^8^. The equivalent mutation in Ace1^12^, Ace2^12,14^, and Mac^13^ rhodopsins was used to generate later GEVIs.

The rhodopsin GEVI optical signal is fast and linear^8^, two desirable features for a voltage indicator. However, rhodopsin fluorescence is very dim, requiring intense illumination for imaging^8^. Fusions of fluorescent protein (FP) domains or other bright fluorophores to rhodopsin GEVIs were therefore made to facilitate imaging^11,13,14^. These fusions enable voltage-sensitive electrochromic fluorescence resonance energy transfer (eFRET) from a bright fluorophore to the retinal cofactor within the rhodopsin, which acts as a dark quencher. While the absorbance of the retinal cofactor increases with increasing membrane potential, the emission of the eFRET-coupled fluorophore consequently decreases. All reported eFRET GEVIs therefore have negatively sloped fluorescence-voltage relationships; they are brighter at resting membrane potential and become dimmer during an action potential (negative-going). Probes with positively sloped fluorescence-voltage relationships (positive-going) are expected to exhibit higher signal-to-noise ratios due to lower background fluorescence. Although two VSD GEVIs, FlicR^4^ and Marina^16^, exhibit positive-going signals in neurons, they have significantly slower response kinetics than eFRET GEVIs, making detection of action potentials difficult. Here, we present a general approach to engineer eFRET GEVIs with fast, bright, and positive-going fluorescence signals in response to neuronal action potentials by modification of the natural proton transport pathway within microbial rhodopsins.

Previous work fused the Ace2 rhodopsin from *Acetabularia acetabulum*^17^ to the FP mNeonGreen to produce a negative-going eFRET GEVI that allowed *in vivo* imaging of voltage signals in several model organisms^12^. We recently used the same Ace2 rhodopsin to engineer a negative-going chemigenetic eFRET GEVI called Voltron, which uses a HaloTag protein domain to covalently bind bright and photostable small molecule flurophores^18,19^, extending the duration and number neurons imaged simultaneously *in vivo*^14^. In both of these GEVIs, photocurrent of Ace2 rhodopsin (Fig. 1a) is blocked by mutating the residue that normally functions as the proton acceptor (PA)^20^ (D81N) (Fig. 1a,b), analogous to the Arch D95N mutation described above. This mutation blocks the primary pathway for exchange of protons from the retinal Schiff base, which links retinal to the rhodopsin protein, to outside the cell^20^. Electrophysiology measurements showing transient inward photocurrents with Ace2 D81N (Fig. 1c) and other rhodopsin based GEVIs^21^ suggest that voltage sensitivity in Ace2 D81N and other eFRET GEVIs results from membrane potential changes altering the equilibrium of protonation between the retinal Schiff base, the proton donor (PD) residue^20^, and the cell cytoplasm (Fig 1a,b).

**Fig. 1.**
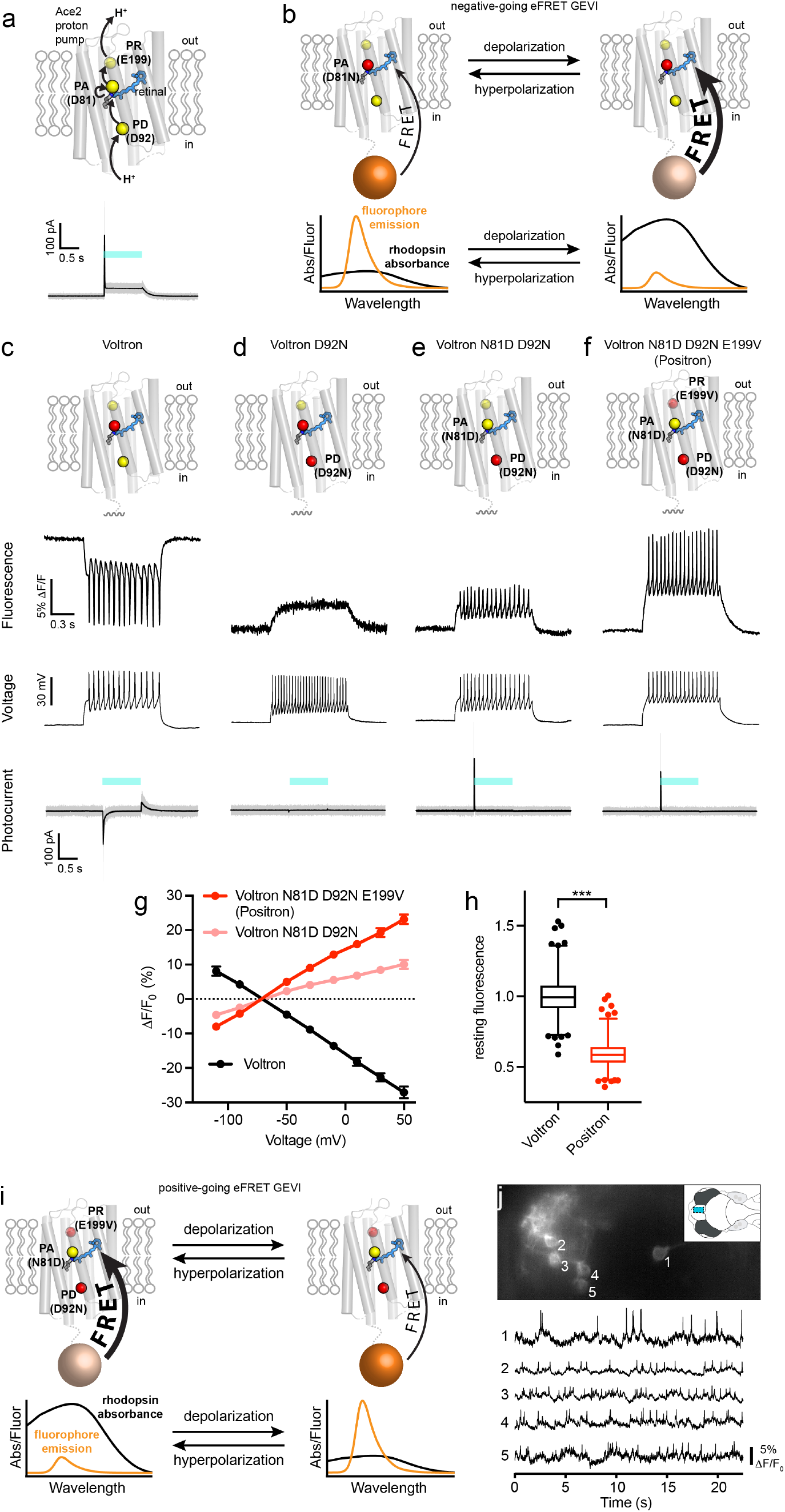
Engineering a positive-going rhodopsin eFRET GEVI. **a**, Schematic (top) showing the hypothetical path of proton transport (arrows) through the Ace2 rhodopsin proton pump. The proton acceptor (PA), proton donor (PD), and proton release (PR) positions are represented as yellow spheres and labeled, retinal is shown as blue sticks. Photocurrent measurements (bottom) for the Ace2 rhodopsin proton pump. Steady state photocurrent = 46 ± 11 pA (mean ± std, N = 4 cells). **b**, Schematic representing the proposed mechanism of negative-going rhodopsin eFRET GEVIs. **c-f**, Schematic of amino acid substitutions (top), simultaneous fluorescence imaging (second row) and whole-cell patch clamp membrane voltage measurements (third row), as well as photocurrent measurements (bottom) from rat hippocampal neurons in culture expressing Voltron, Voltron D92N, Voltron D92N N81D, and Positron labeled with JF_525_. Blue bar denotes time of light illumination ((508 nm – 522 nm) at an irradiance of 70 mW/mm^2^) for photocurrent measurements. Steady state photocurrents for all variants are negligible (-0.7 ± 3 pA, –0.1 ± 1 pA, 0.0 ± 1.5 pA, 0.1 ± 1 pA (mean ± std, N = 5-7 cells) respectively). **g**, Graph of fluorescence vs. membrane voltage for Voltron, Voltron D92N N81D, and Positron expressed in voltage-clamped rat hippocampal neurons in culture. **h**, Box and whisker plot showing the fluorescence of Voltron and Positron at resting membrane potential in neuron culture. Whiskers represent 1^st^-99^th^ percentile. Outliers shown as dots. N = 633 cells and 686 cells. **i**, Schematic representing the proposed mechanism of positive-going rhodopsin eFRET GEVIs. **j**, Imaging of voltage signals from five neurons in the forebrain of live larval zebrafish. (top) Imaged field of view showing fluorescence from five individual neurons expressing Positron and labeled with JF_525_. Inset shows an overview of the larval zebrafish brain and the imaged area (blue rectangle). (bottom) Fluorescence of the five visible neurons over time.

We hypothesized that we could alter the local electrochemical potential of protons on the retinal Schiff base and instead establish a protonation equilibrium with the outside of the cell, which would cause the rhodopsin absorbance and eFRET fluorescence to exhibit the opposite response to membrane potential change. Blocking access of protons to the retinal Schiff base from the cell cytoplasm should enable preferential exchange of protons with the outside of the cell. This hypothesis is supported by a recent report that describes a natural light-driven inward proton pump showing the capacity of microbial rhodopsins to accept protons from outside the cell^22^. To block access of protons from the cytoplasmic side, we substituted the amino acid at the PD position of Voltron for a neutral residue (D92N). As expected, this substitution led to a block of the transient inward photocurrent of Voltron (Fig 1d). Importantly, Voltron D92N (as well as other substitutions to neutral residues at the PD position (Supplementary Fig. 1)) showed a positive-going fluorescence signal with membrane depolarization, but with slow kinetics that made it incapable of following neuronal action potentials (Fig 1d).

We reasoned that the proton pathway between the retinal Schiff base of Voltron D92N and the exterior of the cell was inefficient, resulting in the observed slow kinetics. To improve the efficiency of proton movement towards the outside of the cell, we substituted the amino acid at the PA position for a negatively charged aspartate (N81D), as was present in the original Ace2 rhodopsin sequence. This resulted in an indicator (Voltron N81D D92N) that had sufficient response speed to track action potentials in neurons (Fig. 1e). Critically, Voltron N81D D92N exhibited no steady-state photocurrent (Fig. 1e), showing that it can function as a GEVI without pumping protons across the membrane. Voltron N81D D92N had a transient outward photocurrent (Fig. 1e), confirming that the Schiff base proton was now in equilibrium with the outside of the cell. Although Voltron N81D D92N was suitable for monitoring neuronal action potentials with positive-going fluorescence changes, it showed only ~40% of the fluorescence change of the negative-going Voltron (Fig. 1e, g).

To improve the sensitivity of Voltron N81D D92N we focused on the rest of the proton transport pathway of the Ace2 rhodopsin. We reasoned that proton release (PR) residues^20^ near the outside of the cell should play a critical role in proton movement towards the cell exterior and possibly modulate the protonation equilibrium of the retinal Schiff base. Saturation mutagenesis of the two PR sites (positions 189 and 199 in Ace2) led to a variant with the substitution E199V that had ~2 fold improved dynamic range over Voltron N81D D92N (Fig. 1f,g and Supplementary Fig. 2). We decided to name Voltron N81D D92N E199V “Positron” due to its positive-going response to membrane depolarization. Compared with Voltron, Positron bound to the fluorescent dye JF_525_ had similar fluorescence response to voltage steps, but with a positive fluorescence-voltage slope (Fig. 1g and Supplementary Fig. 3). With sub-millisecond on and off time constants (Supplementary Fig. 4 and Supplementary Table 1), Positron clearly reported action potentials in neurons (Fig. 1f) with sensitivity equivalent to that of Voltron. We observed that Positron was 41% dimmer than Voltron at resting membrane potential in neurons (Fig. 1h), consistent with the hypothesis that at rest, the retinal Schiff base of Positron is more protonated and has a higher absorption, resulting in more eFRET and less dye fluorescence emission. When the membrane depolarizes, the proton is driven toward the outside of the cell, resulting in lower retinal absorption, less quenching of dye fluorescence, and a positive-going fluorescence response (Fig. 1i).

We previously showed that Voltron was suitable for imaging voltage signals in the brains of live animals such as fruit flies, zebrafish, and mice^14^. To confirm that Positron allows for *in vivo* imaging, we recorded optical voltage signals from five neurons simultaneously in the forebrain of a larval zebrafish expressing Positron and labeled with JF_525_ using a widefield fluorescence microscope (Fig. 1j). We observed independent spiking and subthreshold signals in each of the neurons (Fig. 1j) with fluorescence changes and signal-to-noise comparable to those observed with Voltron imaged similarly in the same preparation (compare Fig. 1j with Fig. S34 of ref. 14).

To demonstrate the generality of our approach to generate eFRET GEVIs with positive-going fluorescence response, we explored different reporter fluorophores and rhodopsin domains. Positron showed sensitive fluorescence response when labeled with a yellow dye (JF_525_) or a red dye (JF_585_) (Fig. 2a). We also showed that the HaloTag could be exchanged for either green or red FP domains (Fig. 2b). We created positive-going versions of the Ace2N-mNeon^12^ and VARNAM^23^ eFRET GEVIs capable of sensitively reporting action potentials in neurons with positive-going signals (Fig. 2b and Supplementary Figs. 5 and 6). Next, we substituted the rhodopsin domain with Ace1, Mac, and Arch rhodopsins bearing mutations analogous to the three mutations we introduced to create Positron (Fig. 2c and Supplementary Figs. 7-12). Despite only low amino acid sequence identity (27-54%) between these rhodopsins (Supplementary Fig. 10), each of these permutations resulted in a GEVI with positive-going fluorescence changes capable of following action potentials when we patched neurons in culture and injected current (Fig. 2c and Supplementary Fig. 12). Importantly, all three rhodopsins bearing mutations analogous to Positron showed no steady-state photocurrent (Supplementary Figs. 11 and 12).

**Fig. 2.**
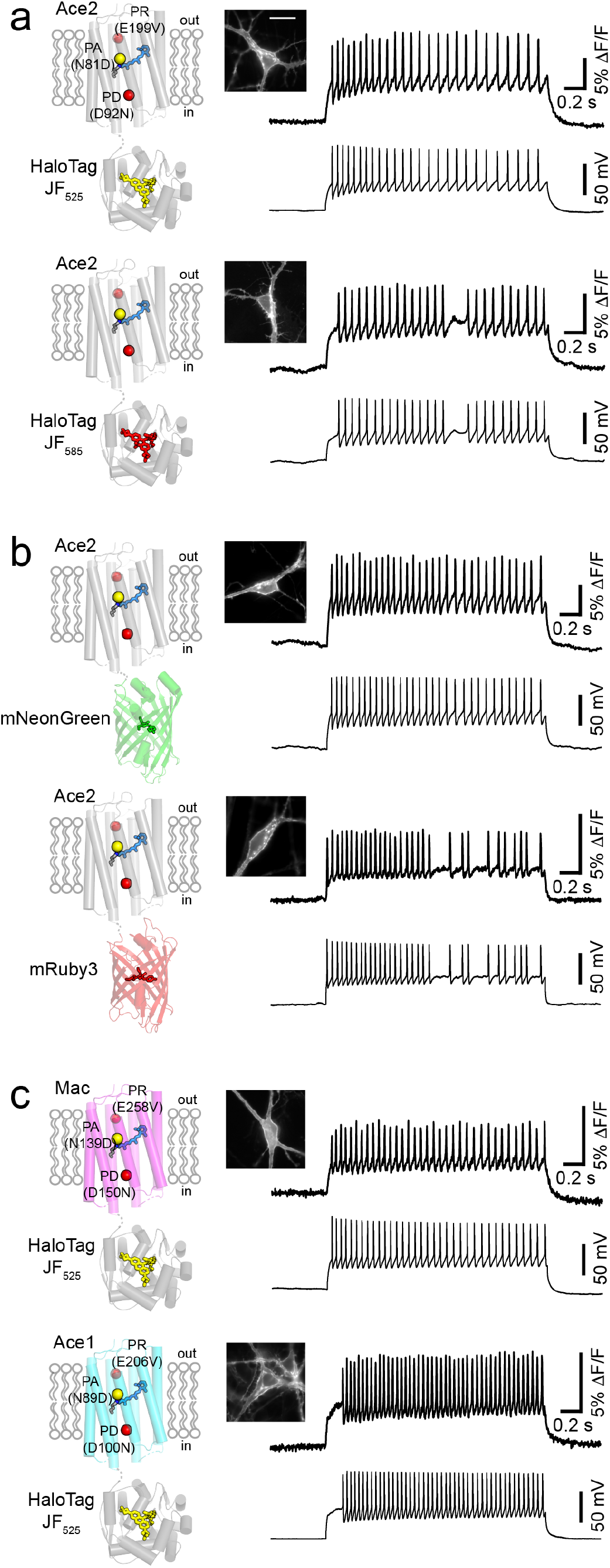
Generality of the approach to engineer positive-going eFRET GEVIs. Each panel shows a schematic of the voltage indicator construct (left), a fluorescence image of a neuron in culture expressing this voltage indicator (center), and simultaneous recording of fluorescence (right top) and membrane potential (right bottom) in response to current injection. Fluorescence image scale bar = 20 μm **a**, Positron labeled with two different colors of fluorescent dye, JF_525_ (top) and JF_585_ (bottom). **b**, Ace2 rhodopsin bearing the signal-inverting mutations described and fused to the different color FP domains mNeonGreen (top) and mRuby3 (bottom). **c**, Different rhodopsin domains (Mac, top and Ace1, bottom) bearing mutations analogous to the described signal-inverting mutations described, fused to HaloTag, and labeled with JF_525_.

Positron is the first eFRET GEVI to run in reverse, having lower fluorescence at resting membrane potentials, positive-going signals in response to action potentials, and sensitivity equal to that of state-of-the-art eFRET GEVIs. We achieved this by rational mutation of the PD, PA, and PR sites within the characterized proton transport pathway of microbial rhodopsins and found that the effect of these substitutions can be generalized to other rhodopsins. This work provides further mechanistic insight into the class of rhodopsin eFRET GEVIs and has potential advantages for lower resting fluorescence background and improved signal detectability in densely labeled samples.

## Acknowledgements

We thank the Janelia Cell Culture, Vivarium, Molecular Biology, and Virus Production facilities for assistance. Specifically, we would like to thank Deepika Walpita, Kim Ritola, and Jordan Towne. Funding was provided by the Howard Hughes Medical Institute. We thank Luke Lavis and Brett Mensh for valuable discussions about the manuscript. We thank the Lavis lab for providing Janelia Fluor dyes. We thank the GENIE project team for contributions to test voltage indicator prototypes.

## Author contributions

A.S.A and E.R.S designed experiments. A.S.A, R.V, A.W, M.K, D.S.K performed and analyzed experiments. A.S.A and E.R.S wrote the manuscript.

## Supplementary Materials

### Methods

#### Reagent availability

Positron plasmids have been deposited at Addgene (www.addgene.org) as follows: pCAG-Positron (plasmid# 129253), pCAG-Positron-ST (plasmid# 129254), pCAG-Ace1_Q89D_D100N_E206V-HaloTag (plasmid# 129255), pCAG-Ace1_Q89D_D100N_E206V-HaloTag-ST (plasmid# 129260), pCAG-Ace2_D92N_E199V-mNeon (plasmid# 129256), pCAG-Ace2_D92N_E199V-mNeon-ST (plasmid# 129261), pCAG-Ace2_D92N_E199V-mRuby3 (plasmid# 129257), pCAG-Ace2_D92N_E199V-mRuby3-ST (plasmid# 129262), pCAG-QuasAr3_Q95D_H106N_E214V-HaloTag (plasmid# 129258), pCAG-QuasAr3_Q95D_H106N_E214V-HaloTag-ST (plasmid# 129263), pCAG-Mac_Q139D_D150N_E258V-HaloTag (plasmid# 129259), pCAG-Mac_Q139D_D150N_E258V-HaloTag-ST (plasmid# 129264), pTol2-Huc-Positron-ST (plasmid# 129265), pT2-Tbait-UAS-Positron-ST (plasmid# 129266), pAAV-hsyn-flex-Positron-ST (plasmid# 129267).

#### Molecular biology

The genes for Ace2 and HaloTag were amplified from a Voltron plasmid^1^. The genes for Ace1, MacQ, QuasAr3, mNeonGreen, mRuby3 were synthesized (Integrated DNA Technologies) with mammalian codon optimization^2–6^. Rhodopsin and fluorescence reporter genes were combined using overlap PCR. Cloning was done by restriction enzyme digest of plasmid backbones, PCR amplification of inserted genes, isothermal assembly, and followed by Sanger sequencing to verify DNA sequences. A soma localization tag^7,8^ (ST) was added using overlap PCR to make soma targeted versions of indicators (e.g. Positron-ST). For expression in primary neuron cultures, sensors were cloned into a pcDNA3.1-CAG plasmid (Invitrogen). For expression in zebrafish, Positron-ST was cloned into the pTol2-HuC vector (for pan-neuronal expression) and into the pT2-Tbait-UAS vector (for Gal4-dependent expression). The DNA and amino acid sequences of Positron, Ace1_Q89D_D100N_E206V-HaloTag, QuasAr3_Q95D_H106N_E214V-HaloTag, Mac_Q139D_D150N_E258V-HaloTag, Ace2_D92N_E199V-mNeonGreen, Ace2_D92N_E199V-mRuby3, are given in Supplementary Figs. 2, and 5-9. Plasmids and maps are available from Addgene.

#### HEK 293 cell culture

Spiking HEK 293 cells^9^ that stably express NaV1.3 and Kir2.1 were used to test the fluorescence response of some Positron mutants using field stimulation. Cells were grown at 37 °C, 5% CO_2_, in Dulbecco’s modified Eagle medium (DMEM) supplemented with 10% FBS, Geneticin (Invitrogen, 10131-027), Penicillin/Streptomycin (Invitrogen, 15140-122), and Puromycin (Invitrogen, A11138-03). All the transfections and assays were done in cells between passage 5 and 15. Plasmids were transfected using calcium phosphate. After transfection, spiking HEK cells were plated on glass-bottom 24-well plates (MaTeK) and cultured for 24 hours before imaging. To label HaloTag-containing constructs, cells were incubated with 100 nM JF-dye HaloTag ligand for 30 minutes.

#### Primary neuron cell culture

All procedures involving animals were conducted in accordance with protocols approved by the Howard Hughes Medical Institute (HHMI) Janelia Research Campus Institutional Animal Care and Use Committee and Institutional Biosafety Committee. Hippocampal neurons extracted from P0 to 1 Sprague-Dawley rat pups were transfected with pcDNA3.1-CAG plasmids of the various indicators by electroporation (Lonza, P3 Primary Cell 4D-Nucleofector X kit) according to the manufacturer’s instruction. After transfection, hippocampal neurons were plated onto 24 well glass bottom plates (MaTek) or 35 mm glass-bottom dishes (MaTek) coated with poly-D-lysine (Sigma). Neurons were cultured for 8-12 days in NbActiv4 medium (BrainBits). To label neurons expressing Positron, Voltron, Ace1_Q89D_D100N_E206V-HaloTag, QuasAr3_Q95D_H106N_E214V-HaloTag, and Mac_Q139D_D150N_E258V-HaloTag, cultures were incubated with 100 nM JF HaloTag ligand for 20-30 mins.

#### Microscopy

Wide-field imaging was performed on an inverted Nikon Eclipse Ti2 microscope equipped with a SPECTRA X light engine (Lumencore) and a 40X oil objective (NA = 1.3, Nikon). Fluorescence images were collected using a scientific CMOS camera (ORCA-Flash 4.0, Hamamatsu) and image acquisition was performed using HCImage Live (Hamamtsu). A FITC filter set (475/50 nm (excitation), 540/50 nm (emission), and a 506LP dichroic mirror (FITC-5050A-000; Semrock)) was used to image Ace2_D92N_E199V-mNeonGreen. A custom filter set (510/25 nm (excitation), 545/40 nm (emission), and a 525LP dichroic mirror (Chroma)) was used to image Voltron, Positron, Ace1_Q89D_D100N_E206V-HaloTag, and Mac_Q139D_D150N_E258V-HaloTag labeled with JF_525_. A quad bandpass filter (set number: 89000, Chroma) was used along with the appropriate color band from the SPECTRA X light source to image QuasAr3_Q95D_H106N_E214V-HaloTag labeled with JF_549_, Positron labeled with JF_585_, and Ace2_D92N_E199V-mRuby3. For time-lapse imaging during field stimulation or simultaneous electrophysiology measurements, neurons were imaged at 400 − 3200 Hz depending on the experiment. All HEK cell and neuron culture imaging was performed in imaging buffer containing the following (in mM): 145 NaCl, 2.5 KCl, 10 glucose, 10 HEPES, pH 7.4, 2 CaCl_2_, 1 MgCl_2_.

To compare brightness of Positron and Voltron at resting membrane potential, we fused mTagBFP2^10^ to the C-terminus of Positron and Voltron to make pCAG-Positron-mTagBFP2 and pCAG-Voltron-mTagBFP2 plasmids. Fluorescence images were acquired from 10 different wells across three independent transfections for each construct in primary hippocampal neurons. To label Voltron-mTagBFP2 and Positron-mTagBFP2 expressing neurons, cultures were incubated with 100 nM JF_525_ HaloTag ligand for 25 minutes, then washed twice with imaging buffer, incubating the cells for 5 minutes during the second wash. An EBFP2 filter set (405/20 nm (excitation), 460/50 nm (emission), and a 425LP dichroic mirror (49021; Chroma)) was used to image the TagBFP2 channel, and a custom filter set (510/25 nm (excitation), 545/40 nm (emission), and a 525LP dichroic mirror (Chroma)) was used to image the JF_525_ channel in Voltron-mTagBFP2 and Positron-mTagBF2 constructs. The ratio of the TagBFP2 channel and JF_525_ channel was calculated using ImageJ software.

#### Field stimulation in spiking HEK cells or primary neuron culture

A stimulus isolator (A385, World Precision Instruments) with platinum wires was used to deliver field stimuli (50V, 1 ms) to elicit HEK cell spiking or action potentials in cultured neurons as described previously^11^. The stimulation was controlled using Wavesurfer and timing was synchronized with fluorescence acquisition using Wavesurfer and a National Instruments PCIe-6353 board.

#### Electrophysiology in primary neuron culture

Filamented glass micropipettes (Sutter Instruments) were pulled to a tip resistance of 4 – 6 MΩ. Pipettes were positioned with a MPC200 manipulator (Sutter Instruments). Whole cell voltage clamp and current clamp recordings were acquired using an EPC800 amplifier (HEKA), filtered at 10 kHz with the internal Bessel filter, and digitized using a National Instruments PCIe-6353 acquisition board at 20 kHz. Data were acquired from cells with access resistance < 25 MΩ. WaveSurfer software was used to generate the various analog and digital waveforms to control the amplifier, camera, light source, and record voltage and current traces.

All electrophysiology measurements were performed in imaging buffer: 145 NaCl, 2.5 KCl, 10 glucose, 10 HEPES, pH 7.4, 2 CaCl_2_, 1 MgCl_2_, adjusted to 310 mOsm with sucrose. Internal solution for current clamp recordings contained the following (in mM): 130 potassium methanesulfonate, 10 HEPES, 5 NaCl, 1 MgCl_2_, 1 Mg-ATP, 0.4 Na-GTP, 14 Tris-phosphocreatine, adjusted to pH 7.3 with KOH, and adjusted to 300 mOsm with sucrose. For current-clamp recordings to generate action potentials, current was injected (20 – 200 pA for 1-2 s) and voltage was monitored.

For fluorescence step responses of Positron to voltage clamp measurements, 500 nM TTX was added to the imaging buffer to block sodium channels. Synaptic blockers (10 μM CNQX, 10 μM CPP, 10 μM GABAZINE, and 1 mM MCPG) were added to block ionotropic glutamate, GABA, and metabotropic glutamate receptors^11^. Internal solution for fluorescence step responses of Positron to voltage clamp recordings contained the following (in mM): 115 cesium methanesulfonate, 10 HEPES, 5 NaF, 10 EGTA, 15 CsCl, 3.5 Mg-ATP, 3 QX-314, adjusted to pH 7.3 with CsOH, and adjusted to 300 mOsm with sucrose.

For fluorescence voltage curves, cells were held at a potential of –70 mV at the start of each recording and then 1 second voltage steps were applied to step the potential from –110 mV to +50 mV in 20 mV increments. Fluorescence images were acquired at 400 Hz using the same microscope described in the “Microscopy” section above. For determining response speed of indicators, fluorescence images were acquired at 3200 Hz in response to a 100 mV potential step delivered to voltage clamped neurons (from −70 mV to +30 mV). Traces were fit to a double exponential function using MATLAB. All recordings were done at room temperature.

#### Photocurrent measurements

Photocurrents were recorded at room temperature in voltage-clamp mode with a holding potential of −70 mV in response to 1s light pulses. Photocurrents were recorded using an EPC800 amplifier (HEKA), filtered at 10 kHz with an internal Bessel filter, and digitized using a National Instruments PCIe-6353 acquisition board at 20 kHz controlled using WaveSurfer. Light was delivered to the clamped neurons using the same microscope described above. Irradiance at the imaging plane was set to 70 mW/mm^2^ determined with a microscope slide power sensor (S170C, Thorlabs).

#### Imaging in zebrafish

In vivo wide-field voltage imaging in zebrafish was performed using the procedure described previously (ref. 1). Briefly, the Positron-ST indicator was transiently expressed by the injection of pT2-Tbait-UAS-Positron-ST (25 ng/μl DNA with 25 ng/μl Tol2 transposase mRNA in E3 medium) in Tg(elavl3:Gal4-VP16) at 1-2 cell stage. JF_525_ was loaded to the injected fish at three day post-fertilization (dpf) by incubation in 3 μM JF_525_ system water solution for 2 hours. The fish with sparsely labeled forebrain neurons were paralyzed by a-bungarotoxin (1 mg/ml) and mounted in low-melting point agarose. Spontaneous activity of forebrain neurons was imaged using a custom wide-field microscope equipped with a 60x 1.0 NA water immersion objective lens (MRD07620, Nikon), a LED light source (CBT-90-W, Luminous) and a TRITC filter set (TRITC-B-000, Semrock). The images were acquired with sCMOS camera (pco.edge 4.2, PCO) at 400 Hz for 1-2 min. Irradiance at the imaging plane was 28 mW/mm^2^ (S170C, Thorlabs).

---

**Supplementary Figure 1.**
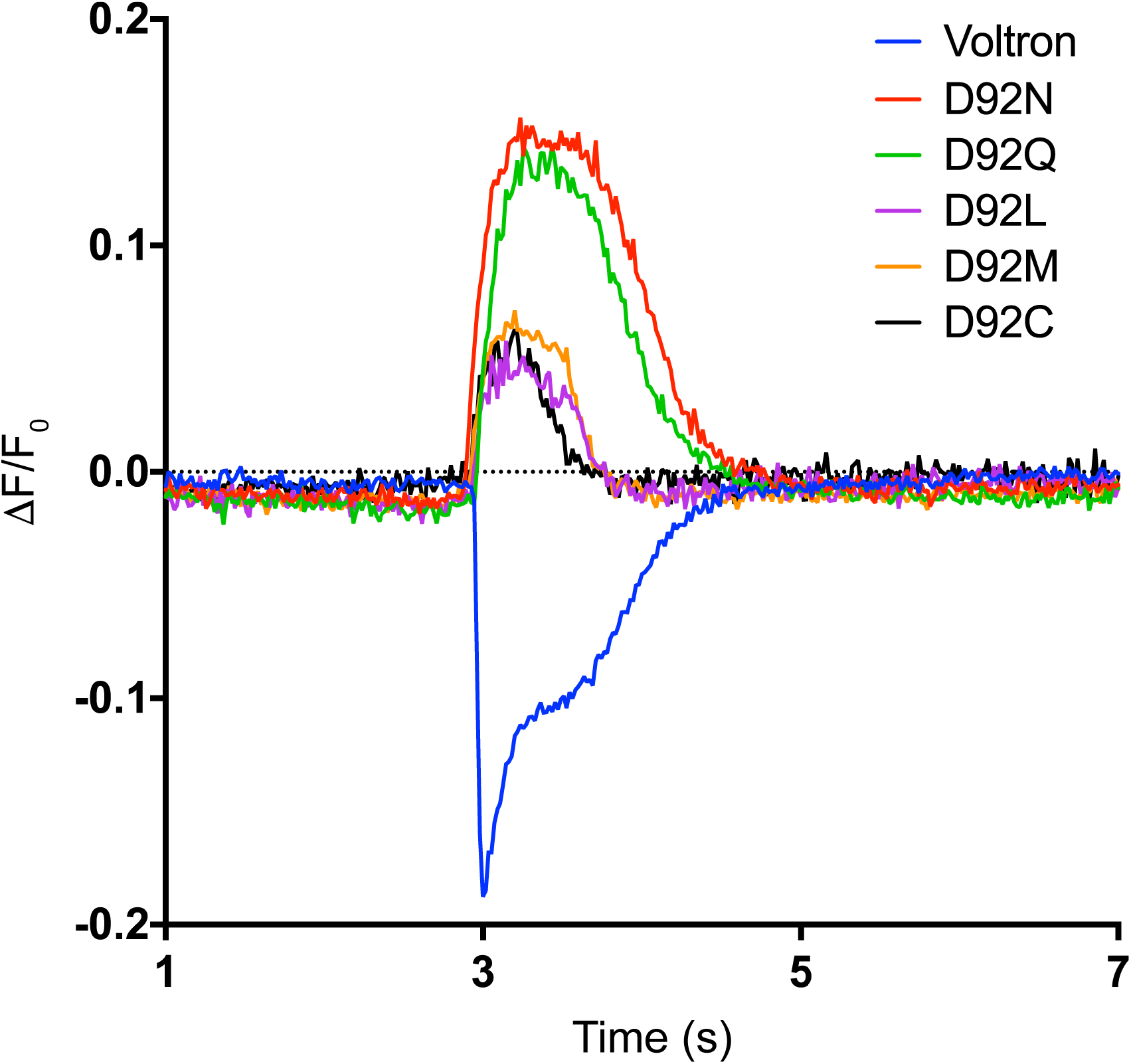
Spiking HEK cell fluorescence traces for Voltron variants with mutations at position D92. Cells were subject to field stimulation at t = 3 seconds. Saturation mutagenesis at the proton donor position of Ace2 (D92) results in variants with a positive fluorescence change in response to membrane depolarization. All Voltron variants were labeled with JF_525_.

**Supplementary Figure 2.**
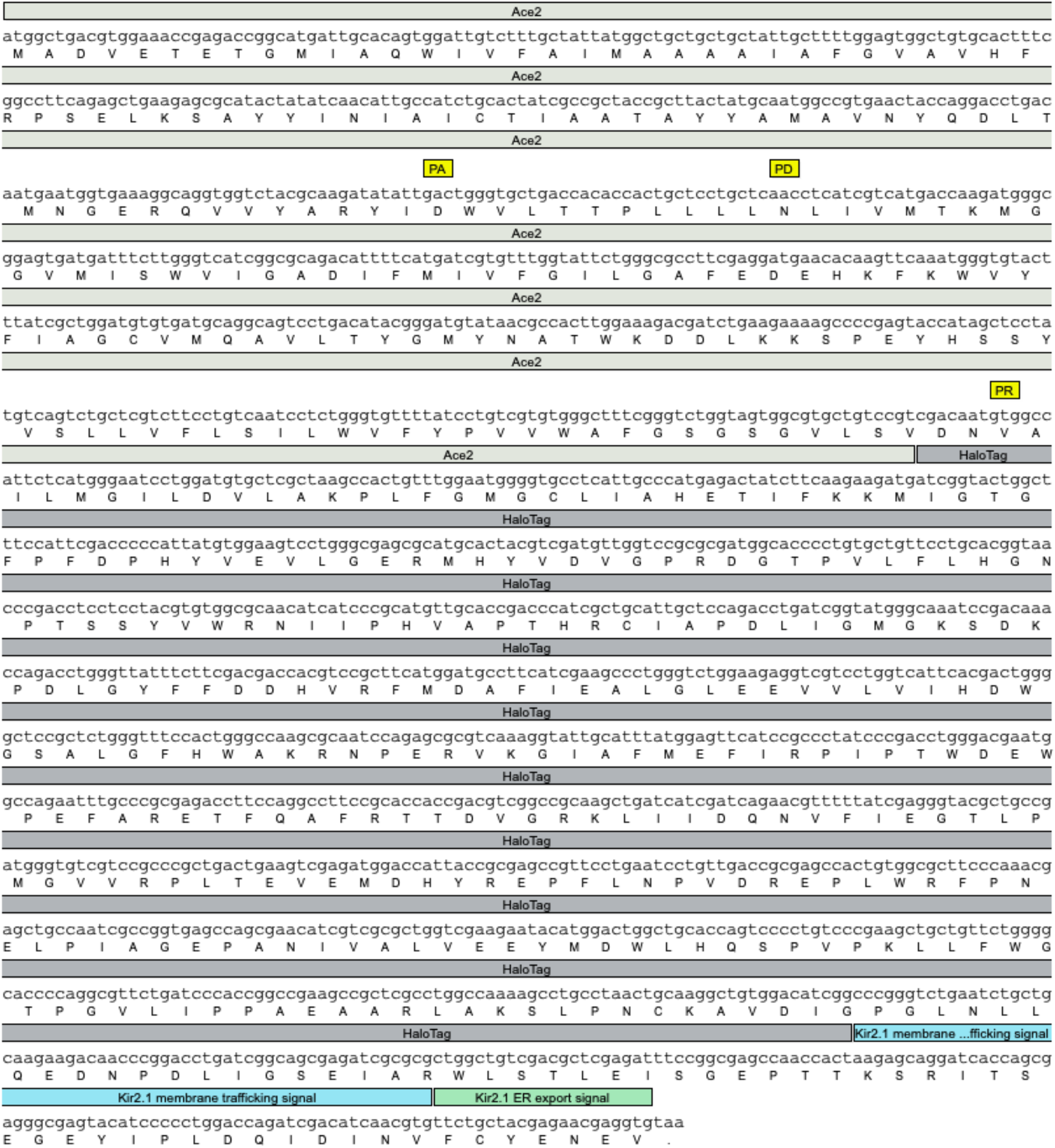
DNA and amino acid sequence of Positron with sequence features annotated. Note the three point mutations that are responsible for the positive fluorescence voltage slope annotated as PA (proton acceptor position), PD (proton donor position), and PR (proton release position).

**Supplementary Figure 3.**
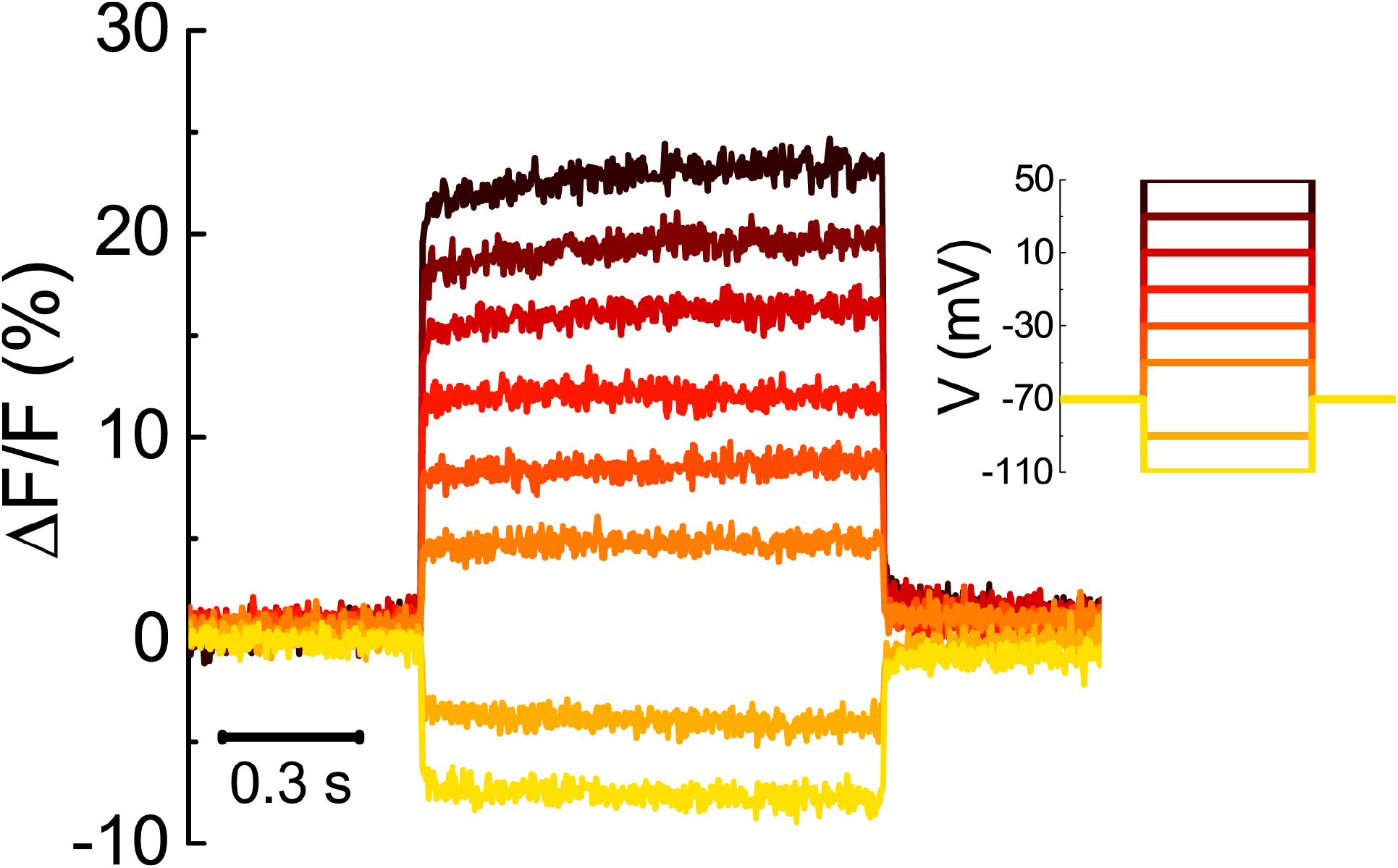
Fluorescence traces of Positron labeled with JF_525_ in response to a series of voltage steps (from −110 mV to +50 mV in 20 mV increments). Image acquisition rate = 400 Hz. For complete fluorescence vs voltage plots of Positron, Voltron_N81D_D92N, and Voltron see Figure 1g.

**Supplementary Figure 4.**
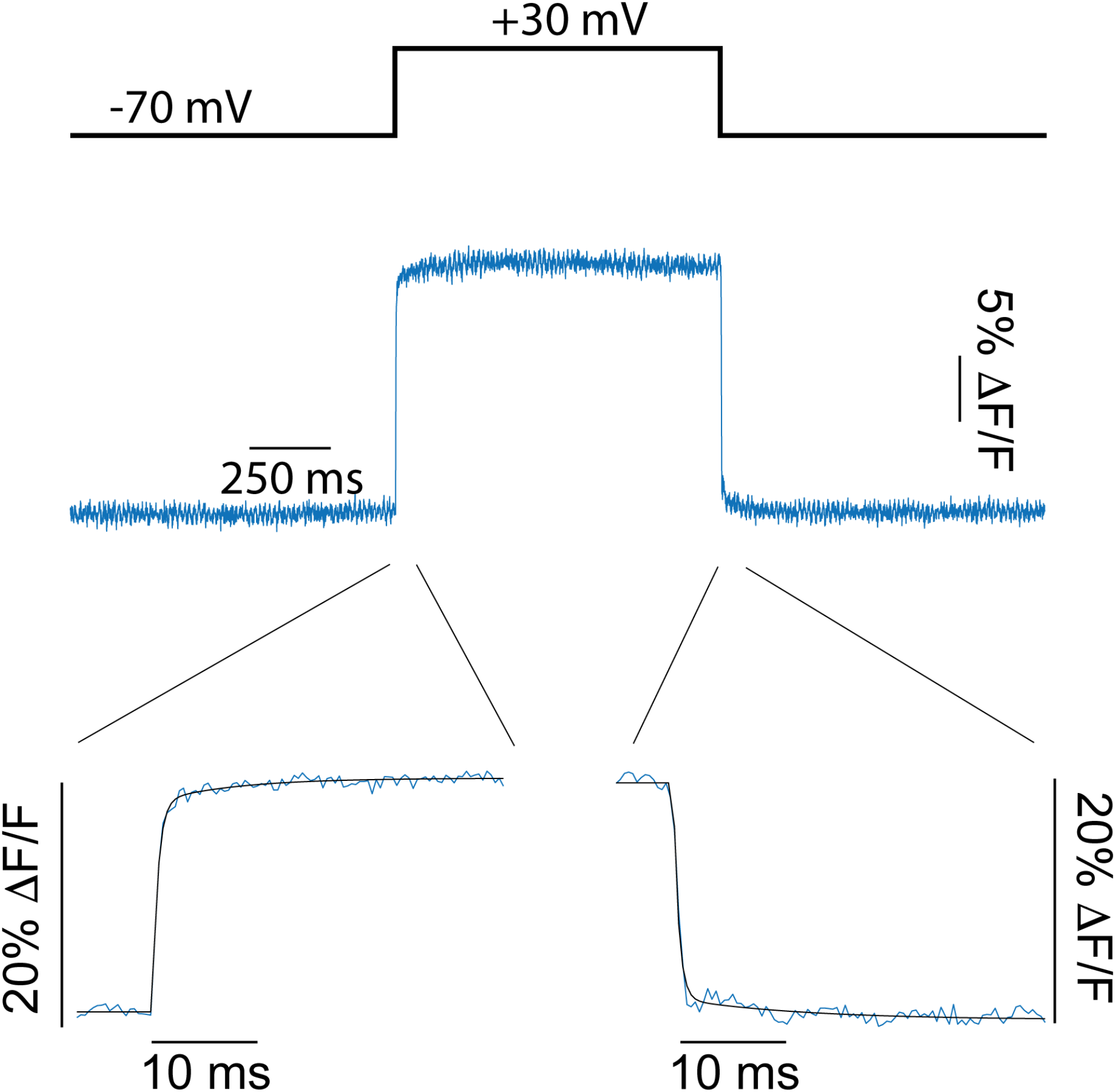
Fluorescence response of Positron labeled with JF_525_ in response to a 100 mV potential step delivered to a voltage clamped neuron. Insets: Zoom in on Positron fluorescence response to membrane depolarization (from −70 mV to +30 mV), and repolarization (from +30 mV to −70 mV). Solid black line is fit of rise and decay kinetics to a double exponential function. Image acquisition rate 3.2 kHz. For full kinetics data, see Supplementary Table 1.

**Supplementary Figure 5.**
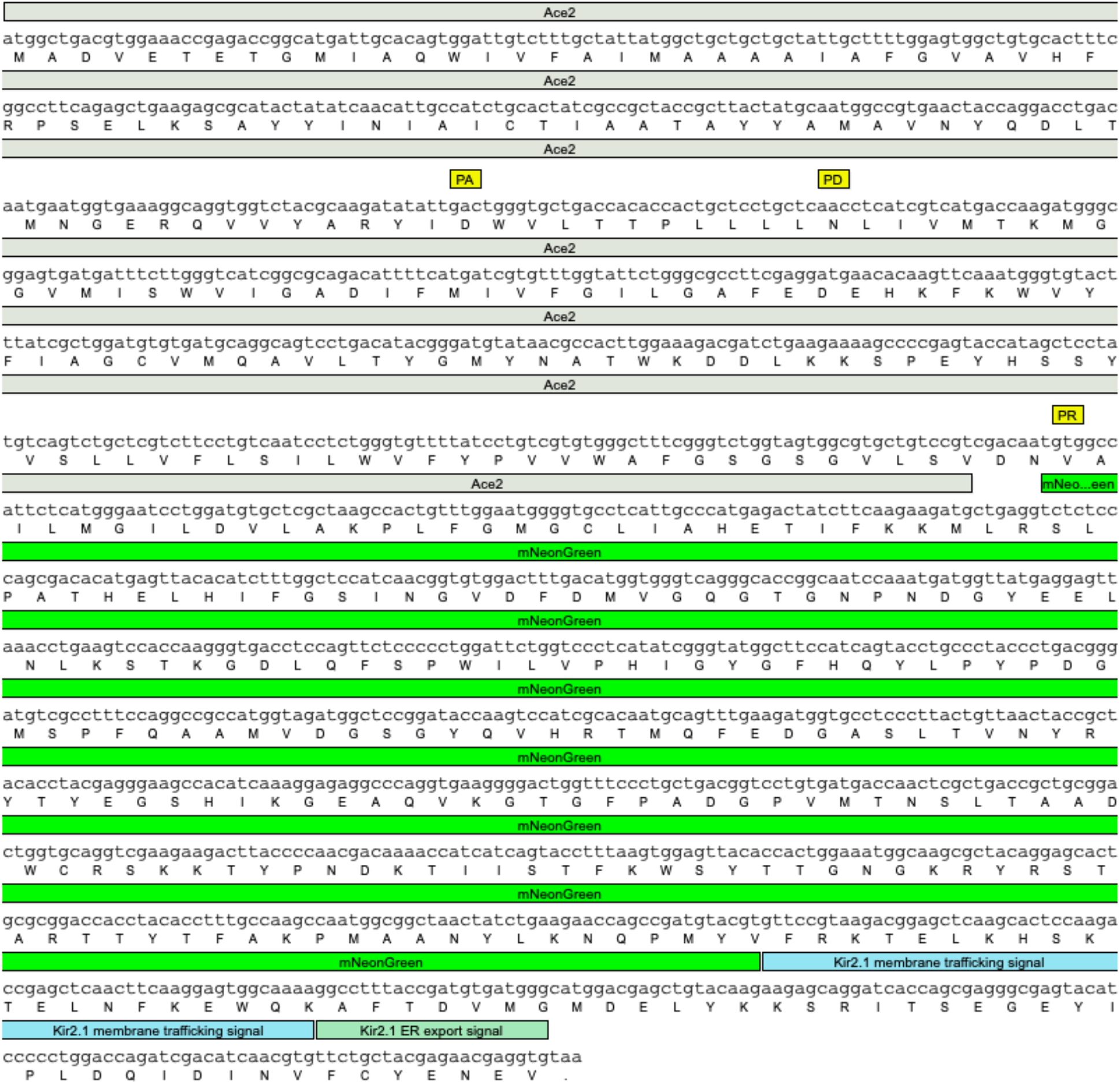
DNA and amino acid sequence of Ace2_D92N_E199V-mNeonGreen with sequence features annotated. Note the three point mutations that are responsible for the positive fluorescence voltage slope annotated as PA (proton acceptor position), PD (proton donor position), and PR (proton release position).

**Supplementary Figure 6.**
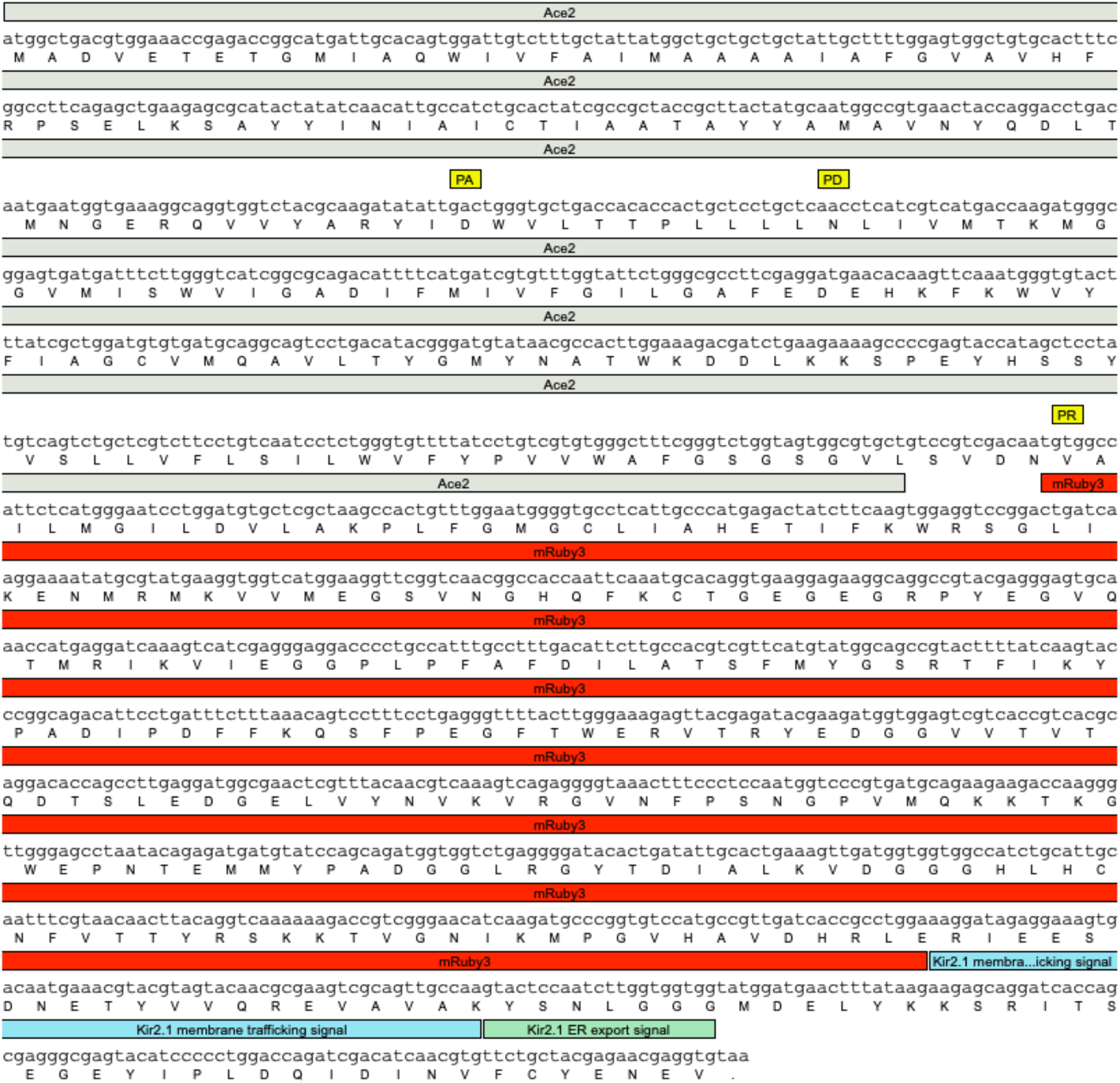
DNA and amino acid sequence of Ace2_D92N_E199V-mRuby3 with sequence features annotated. Note the three point mutations that are responsible for the positive fluorescence voltage slope annotated as PA (proton acceptor position), PD (proton donor position), and PR (proton release position).

**Supplementary Figure 7.**
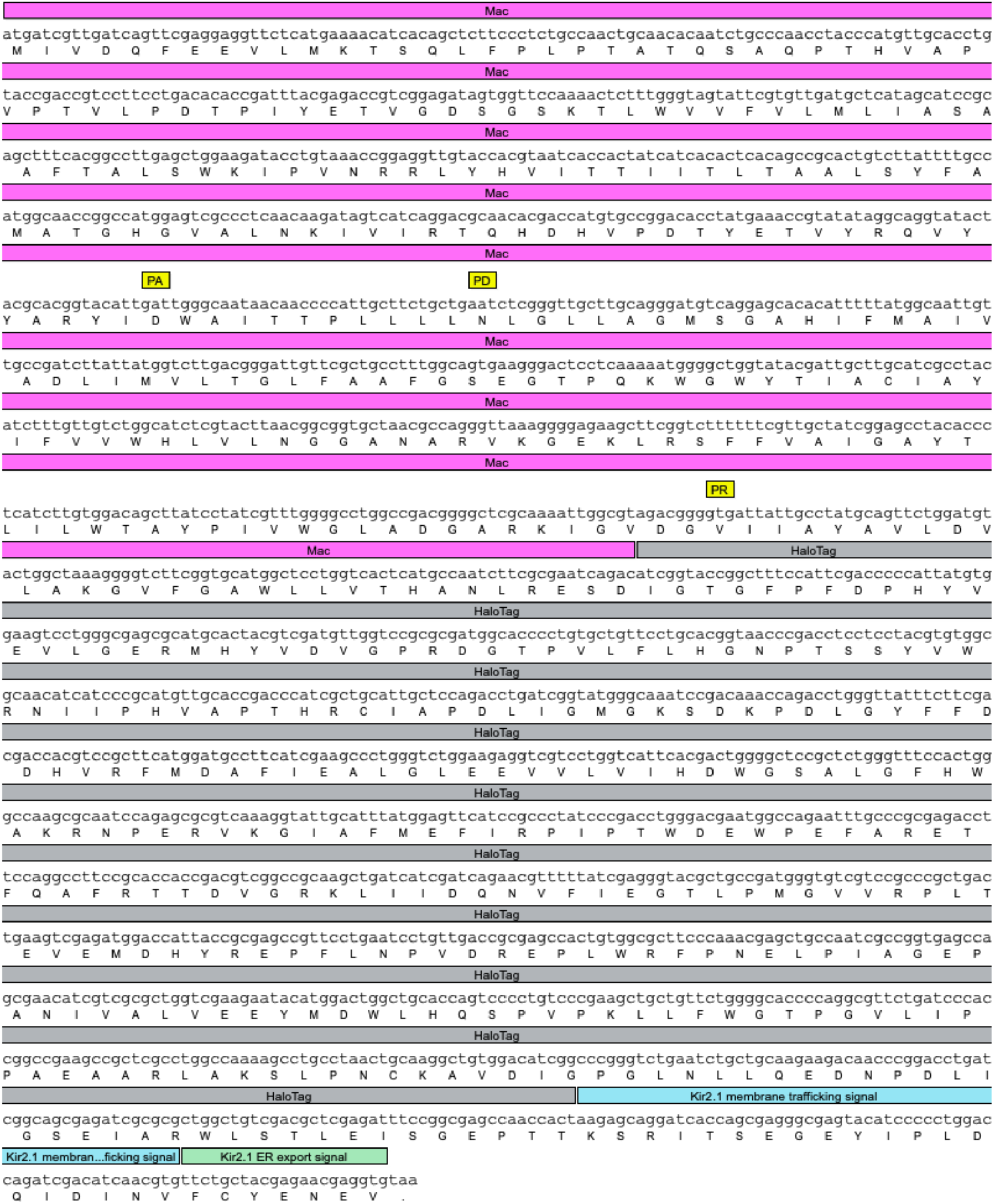
DNA and amino acid sequence of Mac_Q139D_D150N_E258V-HaloTag with sequence features annotated. Note the three point mutations that are responsible for the positive fluorescence voltage slope annotated as PA (proton acceptor position), PD (proton donor position), and PR (proton release position).

**Supplementary Figure 8.**
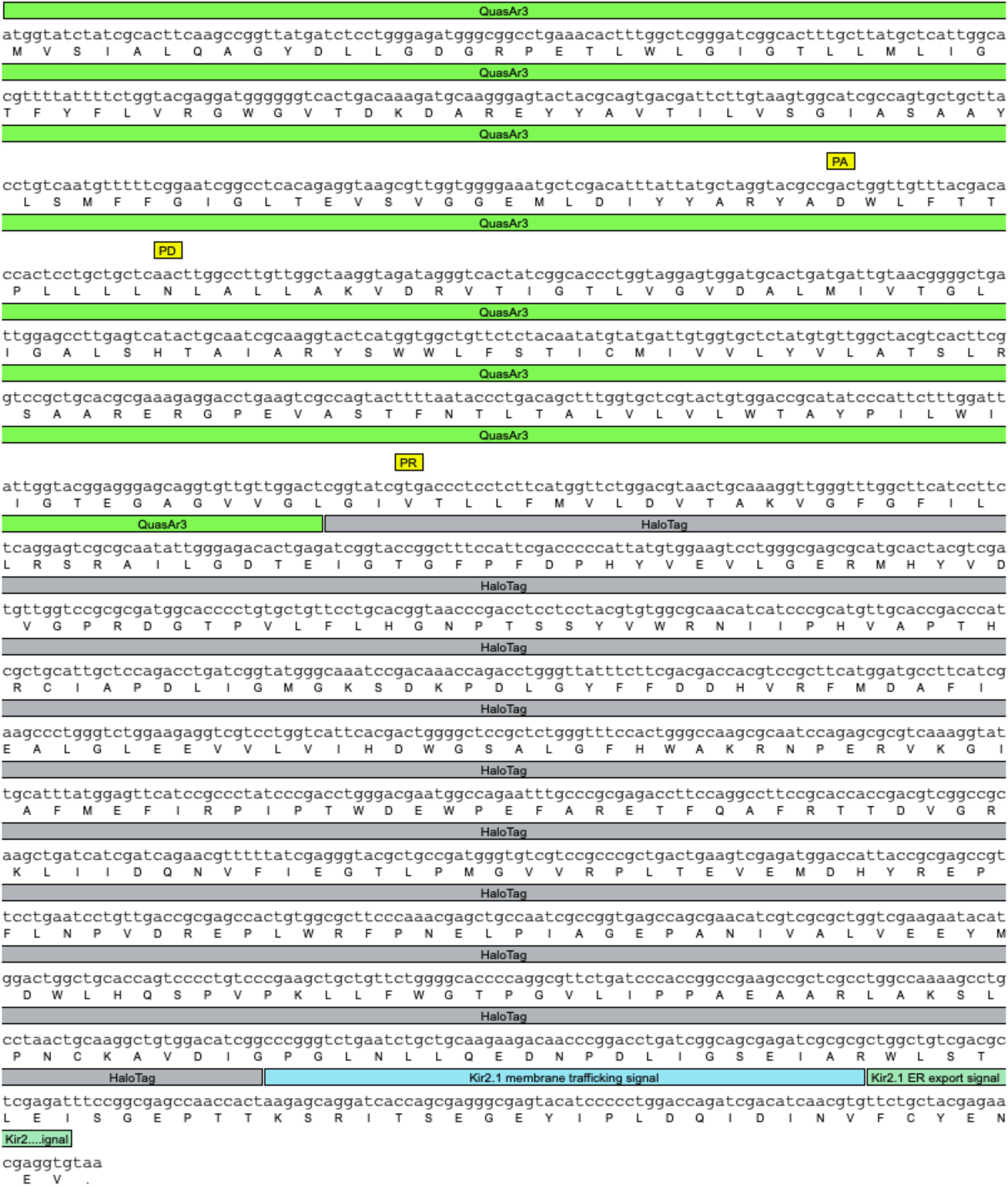
DNA and amino acid sequence of QuasAr3_Q95D_H106N_E214V-HaloTag with sequence features annotated. Note the three point mutations that are responsible for the positive fluorescence voltage slope annotated as PA (proton acceptor position), PD (proton donor position), and PR (proton release position).

**Supplementary Figure 9.**
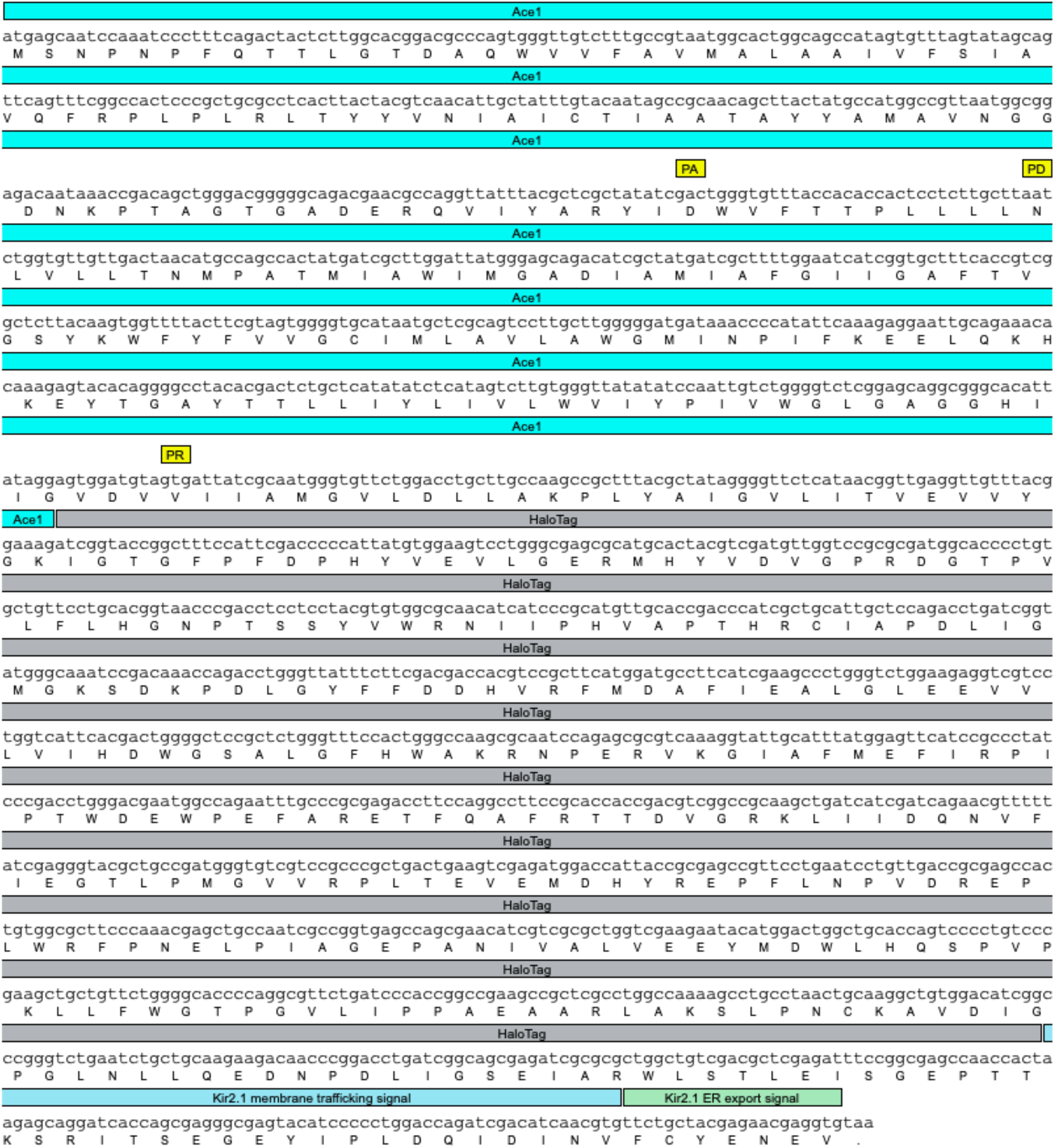
DNA and amino acid sequence of Ace1_Q89D_D100N_E206V-HaloTag with sequence features annotated. Note the three point mutations that are responsible for the positive fluorescence voltage slope annotated as PA (proton acceptor position), PD (proton donor position), and PR (proton release position).

**Supplementary Figure 10.**
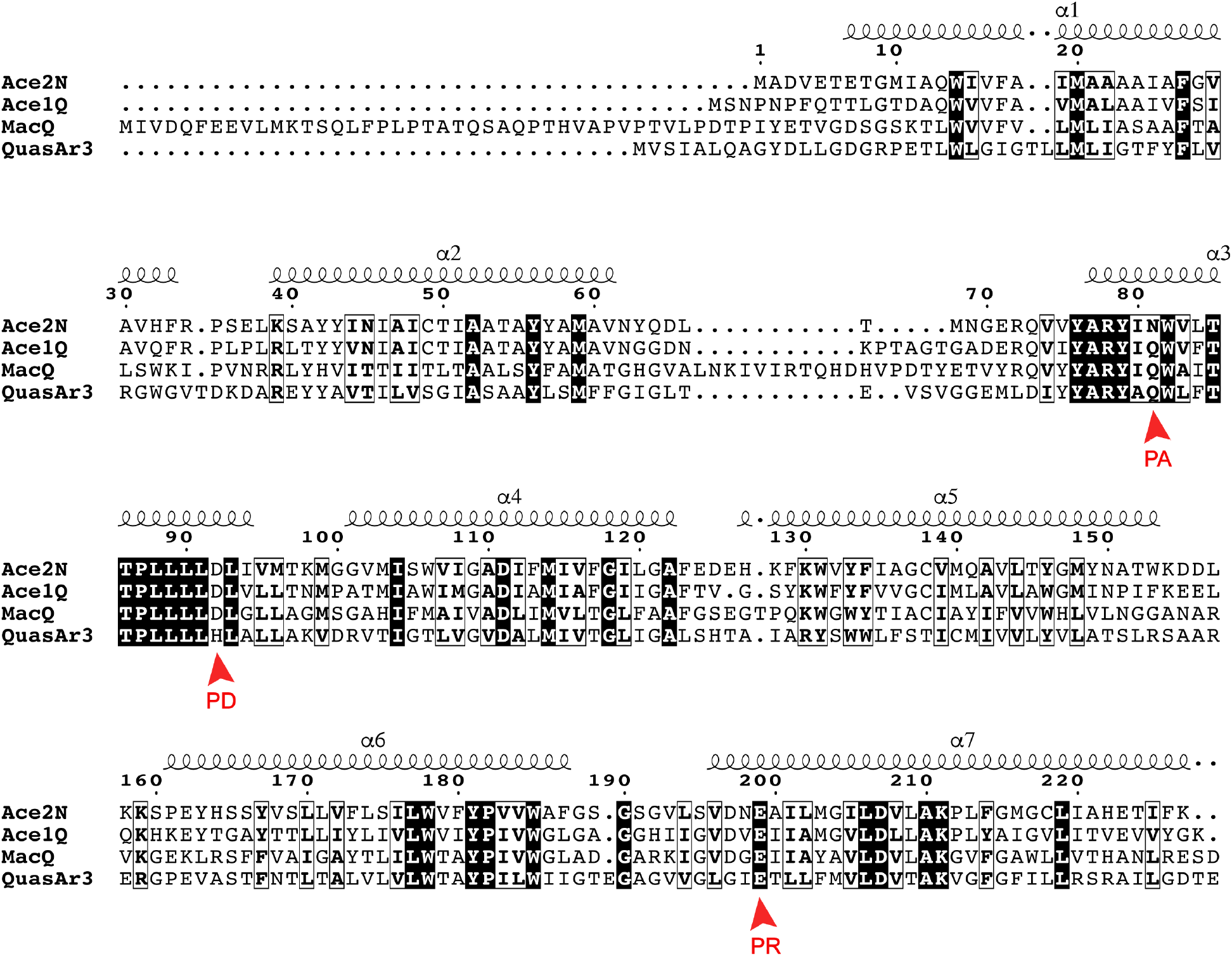
Sequence alignment of the four microbial rhodopsins (Ace2N, Ace1Q, MacQ, and QuasAr3) used to construct eFRET GEVI indicators. The proton acceptor (PA), proton donor (PD), and proton release (PR) residues are labeled with red arrows.

**Supplementary Figure 11.**
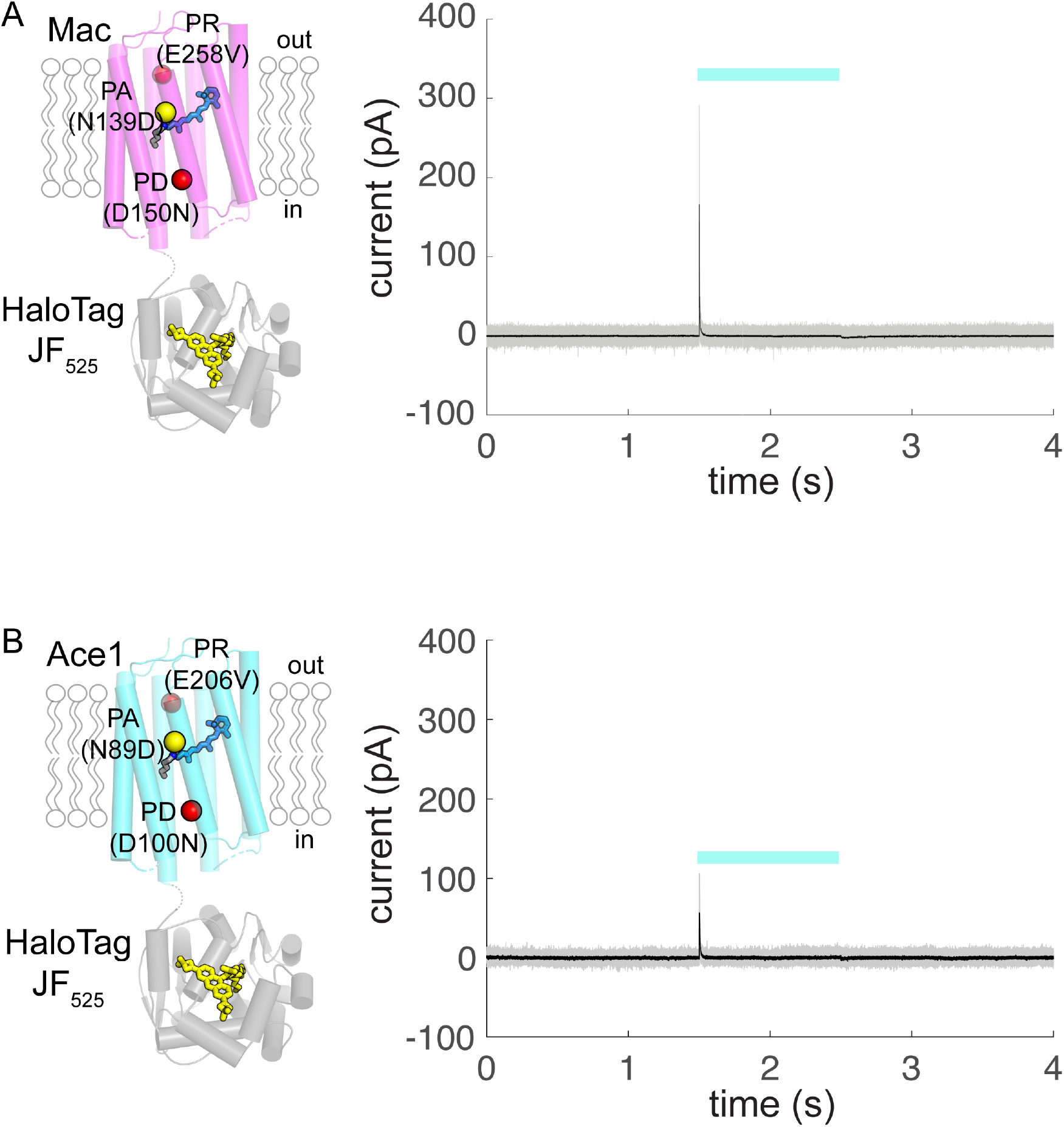
Photocurrent measurements for (A) Mac_Q139D_D150N_E258V-HaloTag and (B) Ace1_Q89D_D100N_E206V-HaloTag. Both constructs showed transient outward currents and no steady state photocurrent (0.0 ± 1.7 pA, and –0.6 ± 1.4 pA (mean ± std) for Mac_Q139D_D150N_E258V-HaloTag (N = 3 cells) and Ace1_Q89D_D100N_E206V-HaloTag (N = 4 cells) respectively). Light used was blue/green (508 nm – 522 nm) at an irradiance of 70 mW/mm^2^ at the specimen plane. Gray traces: individual current recordings. Black trace: Average of individual current recordings.

**Supplementary Figure 12.**
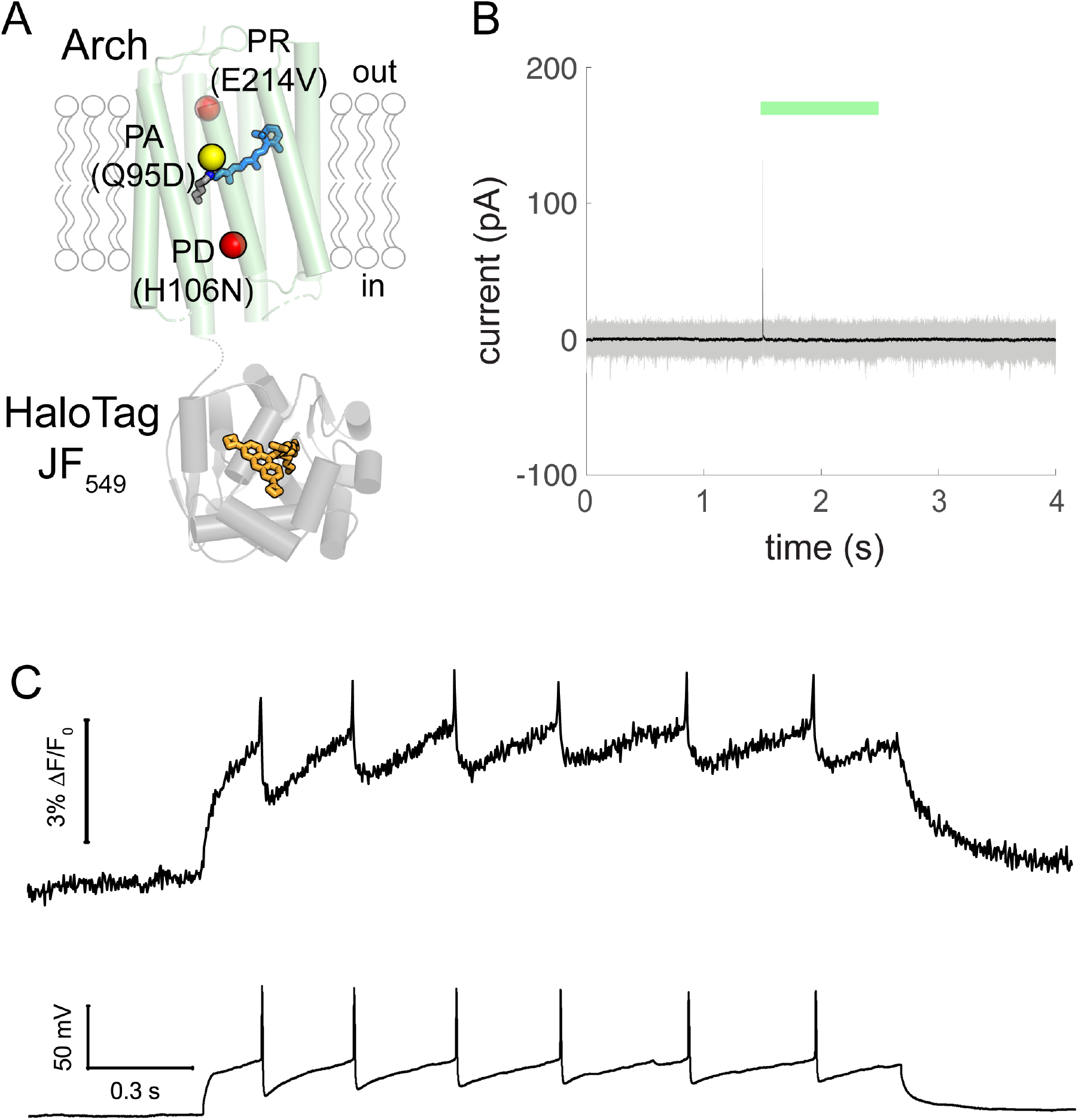
(A) Schematic representation of QuasAr3_Q95D_H106N_E214V-HaloTag labeled with JF_549_ dye. Yellow and red spheres are mutations compared to QuasAr3 sequence. (B) Photocurrent measurements for QuasAr3_Q95D_H106N_E214V-HaloTag. QuasAr3_Q95D_H106N_E214V-HaloTag has a transient outward current and no steady state photocurrent (-0.8 ± 3 pA, (mean ± std) (N = 4 cells)). Light used was green (540 nm – 570 nm) at an irradiance of 70 mW/mm^2^ at the specimen plane. Gray traces: individual current recordings. Black trace: Average of individual current recordings. (C) Simultaneous recording of fluorescence (top) and membrane potential (bottom) in response to current injection of a neuron expressing QuasAr3_Q95D_H106N_E214V-HaloTag and labeled with JF_549_.

**Supplementary Table 1.**
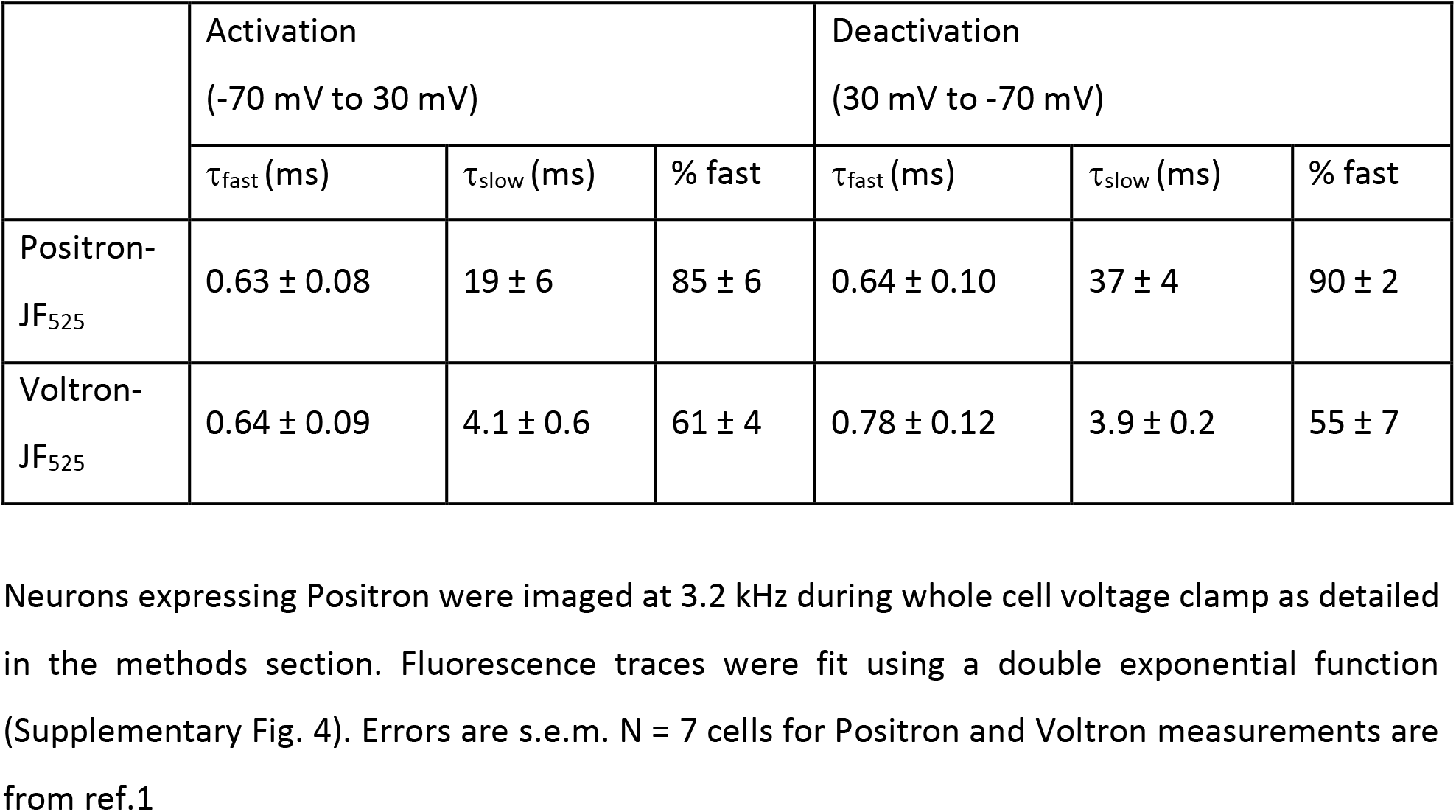
Positron and Voltron kinetics in primary rat neuron cultures

|  | Activation<br>(-70 mV to 30 mV) |  |  | Deactivation<br>(30 mV to -70 mV) |  |  |
| --- | --- | --- | --- | --- | --- | --- |
| | $\tau_{\text{fast}}$ (ms) | $\tau_{\text{slow}}$ (ms) | % fast | $\tau_{\text{fast}}$ (ms) | $\tau_{\text{slow}}$ (ms) | % fast |
| Positron-JF<sub>525</sub> | $0.63 \pm 0.08$ | $19 \pm 6$ | $85 \pm 6$ | $0.64 \pm 0.10$ | $37 \pm 4$ | $90 \pm 2$ |
| Voltron-JF<sub>525</sub> | $0.64 \pm 0.09$ | $4.1 \pm 0.6$ | $61 \pm 4$ | $0.78 \pm 0.12$ | $3.9 \pm 0.2$ | $55 \pm 7$ |

